# Optimal clustering for quantum refinement of biomolecular structures: Q|R#4

**DOI:** 10.1101/2022.11.24.517825

**Authors:** Yaru Wang, Holger Kruse, Nigel W. Moriarty, Mark P. Waller, Pavel V. Afonine, Malgorzata Biczysko

**Affiliations:** International Center for Quantum and Molecular Structures, Physics Department, College of Sciences, Shanghai University, Shanghai 200444, People’s Republic of China; Institute of Biophysics of the Czech Academy of Sciences, Brno, Czech Republic; Molecular Biosciences and Integrated Bioimaging, Lawrence Berkeley National Laboratory, Berkeley, CA 94720, USA; Pending AI Pty Ltd., iAccelerate, Innovation Campus, North Wollongong, 2500, Australia

## Abstract

Quantum refinement (Q|R) of crystallographic or cryo-EM derived structures of biomolecules within the Q|R project aims at using *ab initio* computations instead of library-based chemical restraints. An atomic model refinement requires the calculation of the gradient of the objective function. While it is not a computational bottleneck in classic refinement it is a roadblock if the objective function requires *ab initio* calculations. A solution to this problem adopted in Q|R is to divide the molecular system into manageable parts and do computations for these parts rather than using the whole macromolecule. This work focuses on the validation and optimization of the automatic *divide-and-conquer* procedure developed within the Q|R project. Also, we propose an atomic gradient error score that can be easily examined with common molecular visualization programs. While the tool is designed to work within the Q|R setting the error score can be adapted to similar fragmentation methods. The gradient testing tool presented here allows *a priori* determination of the computationally efficient strategy given available resources for the potentially time-expensive refinement process. The procedure is illustrated using a peptide and small protein models considering different quantum mechanical (QM) methodologies from Hartree-Fock, including basis set and dispersion corrections, to the modern semi-empirical method from the GFN-xTB family. The results obtained provide some general recommendations for the reliable and effective quantum refinement of larger peptides and proteins.

## 1. Introduction

Accurate and reliable structures of bio-macromolecules at an atomistic level have been instrumental in our understanding of biological processes [1, 2, 3], drug design [4] or protein engineering [5]. Atomic model refinement provides an efficient route to obtain high-quality models of such molecules. Refinement takes an atomic model and improves it to optimally match the data that originates from a crystallographic or cryo-EM experiment [6]. However, the limited resolution of the experimental data and low data-to-parameter ratio require the refinement procedure uses *a priori* chemical information. This chemical information is utilized as stereo-chemical restraints or constraints [7, 8] (such as bond lengths, bond angles, torsion angles, and so on) and originates from various sources, such as small molecule databases (Cambridge Structural Database (CSD) [9, 10], Crystallography Open Database (COD) [11]) or the ultrahigh-resolution protein structures (Conformation Dependent Library (CDL) [12, 13, 14]). These restraints are limited to the information available in the databases and fail to easily cope with novel chemical moieties such as new drugs [15, 16, 17, 18] or unusual local arrangements [19, 20, 21]. Also, they do not account for electrostatic effects [19] or local and typically unique to the structure non-covalent interactions such as hydrogen bonds, salt bridges, π-stacking and more [22, 23].

Quantum mechanical (QM) methods have proven to be powerful, predictive tools in protein structure research [24, 25, 26, 27, 28, 29]. One of the examples is our newly developed quantum-based refinement (Q|R) package [30] that combines QM and crystallographic [26] or cryo-EM [27] experimental data. The Q|R software is developed as an open-source module of the Phenix package [31]. It does not require any of the static library-based parameterized restraints (such as Monomer Library; [32, 33]) using QM calculations instead. The existing developments that use quantum mechanical calculations as a source of restraints for macromolecular crystallographic refinement employ methods based on multiscale QM/molecular mechanics (MM) [34, 35], semi-empirical [3] or linear-scaling density functional theory [36]. These tools typically focus on the region of particular interest in the macro-molecule (e.g., ligand or ligand-binding pocket) while neglecting the rest of the model or treat it with the standard classic approach using stereo-chemical restraints. In Q|R based refinements the whole protein structure is treated at the QM or semi-empirical (SQM) level, for example, the HF/6-31G level, employing also London dispersion (D3) [37] and basis set incompleteness (gCP) corrections [38] or by GFN-xTB [39], accounting for the polar environment by means of continuum solvent model (CPCM/COSMO [40] or internal xtb generalized born (GB) solvation model).

QM restraints provide more chemically meaningful bio-macromolecular structures [41]. However, the issue of computational scalability in QM methods [42] is one of the main obstacles to perform reliable and efficient QM calculations in the quantum-based refinement. Although computational algorithms and hardware resources steadily improve, this issue remains [43].

The atomic model refinement requires calculation of an objective function value and its gradients with respect to refinable parameters (e.g., Cartesian coordinates), where the objective function is a weighted sum of stereochemical restraints (classic or quantum) and the experimental data term [6]. To address the scalability issue the whole QM gradient can be computed by one of the *divide-and-conquer* methods [29, 44, 45, 46, 47]. Assuming that the molecules in the local area of the macro-molecular system are hardly affected by the molecules further away, in principle, any large protein can be divided into a group of smaller pieces [44, 48]. In the Q|R project we are using our own partitioning scheme [49] where the macromolecule is divided into *clusters*. A *fragment* contains the *cluster* and its surrounding *buffer* region. This partitioning algorithm is crystallographic symmetry-aware [50]. Since the energies arising from individual clusters cannot be combined into the total energy, the Q|R refinement only uses the energy gradients. The QM energy gradients for all atoms from each of the *fragments* are computed with only the gradients from each *cluster* combined to create the total gradient. This procedure has been already successfully applied to improve several molecular models by quantum refinement in connection with experimental data from X-ray crystallography [49, 50] and cryo-EM [51]. It has been also used in the QM optimization of protein structure [51]. The *divide-and-conquer* procedure can potentially introduce and accumulate errors at the dividing boundaries of the *clusters* due to, for example, insufficient size of the *buffer* region. This problem can be readily alleviated by expanding the *buffe*r region which in turn will increase the time of computations (often by a large margin).

Here we continue our series [30, 49, 50, 51] of publications that document the evolution and progress of the Q|R project. In what follows we describe how *divide-and-conquer* procedure can be optimized to minimize these errors without substantial increase of the computational time, which sets another milestone along the road of the Q|R project.

## 2. Methods

### 2. Model selection and preparation

We use two of the models from the test set of 70 peptide and protein structures employed in previous works [52]. The first model (PDB entry 3ftL, 17 residues, 109 atoms, 1.60 Å resolution) is sufficiently small to allow the detailed analysis of all possible clustering schemes as well as an extensive testing of parameters defining *fragments* for which QM computations are performed. 3ftL is an amyloid peptide with highly ordered and packed aggregates stabilized by intermolecular contacts spanning across symmetry copies [53]. This makes this model a suitable candidate for a validation and analysis of Q|R *divide-and-conquer* procedure. The second model (PDB entry 3q2c, 128 residues, 787 atoms, 2.50 Å resolution) has been selected as a larger and more complex example with different secondary-structure patterns. These models were prepared for test as following. The atomic model and reflection data files have been obtained from the RCSB PDB Database [54] and then refined using *phenix*.*refine* [31, 55] using default settings in order to obtain an optimal starting point. The model was next completed by adding hydrogen atoms using the *qr*.*finalise* [30] tool that is part of the Q|R software suite. The resulting models (3ftL 209 atoms, 3q2c 1611 atoms) have been used for the gradient analysis using Q|R. Additionally, the similar tests have been performed for the models used previously [51].

The crystallographic symmetry-aware *divide-and-conquer* procedure in Q|R [49, 50] includes expansion of the unit cell content in space, which is referred to as *super-sphere*. The *super-sphere* contains the model in question and residues from symmetry copies that fall within a pre-defined distance (R_SS_) from the model. This symmetry expansion distance (R_SS_) is set to 10 Å by default. Once the *super-sphere* is obtained, the model is partitioned into *fragments*, where each fragment is composed of one *cluster* and surrounding *buffer* region. The *buffer* region is needed to account for covalent and non-covalent interactions and should be sufficiently large for accurate QM calculations. If needed, the *buffer* region can be further extended to create a so-called *double-buffer* by adding another layer of residues, interacting with a previously constructed (*single-buffer*) fragment. It should be noted that the R_SS_ distance does not affect division of model into *clusters*, but only influences the size of *buffer region*.

### 2.2 Automated testing for accuracy of QM gradient

Clustered gradient is the gradient for the whole model that is calculated as an accumulation of individual gradients arising from each *cluster*. Therefore, the clustered gradient is an approximation to the exact gradient calculated for the whole model. This is because surrounding atoms interact with the *cluster* through covalent and non-covalent interactions. Inaccuracies of the clustered gradient arise from the two aspects of making a cluster: cutting through covalent bonds (followed by capping of naked atoms at the cut) and limiting the non-covalent interactions. Calculation of the clustered gradient (and therefore its accuracy) is governed by several parameters, such as (i) R_SS_ distance used to define the *super-sphere*, (ii) maximum allowed number of residues in each *cluster*, (iii) size of the *buffer* region surrounding each *cluster* (e.g., *single-* or *double-buffer*). The larger the *super-sphere* and the thicker the *buffer*, the more accurate clustered gradient will be and the more time will be required to calculate it. However, the gradient from the whole model and the clustered gradient can never match exactly due to numerical errors and errors arising from capping. An automated procedure described here aims at obtaining the set of these parameters that minimize the errors in the clustered gradient using the least computational time possible. For this a new command-line tool was added to the Q|R project named *qr*.*gtest*. To evaluate the error in the clustered gradient a reference to compare with (e.g., the exact gradient) is needed. This reference gradient is calculated only once either from the whole *super-sphere* (if computationally possible) or from the clustered gradient obtained with the sufficiently large *buffer* region (if the molecule is too large to use the whole *super-sphere*). In this work we use the gradient calculated from the *super-sphere* as the reference.

There are two options available in the *qr*.*gtest*. One will proceed with the computation of the reference gradient where the model is not divided at all. In that case the QM gradient is computed for the whole *super-sphere*. The second option tests (i) different possibilities of the maximum allowed number of residues in each *cluster*, which define how the model is divided; and (ii) the size of the *buffer* region needed for each cluster. In all cases *qr*.*finalise* is used to cap atoms to assure correct atomic valences at cutting points.

In this work we have tested the effect of the *super-sphere* size on the gradient error by considering the R_SS_ distance between 1 and 30 Å for the computations of QM gradients. Additionally, we analyzed the *fragment* size and composition obtained with *single-* and *double-buffer*.

### 2.3 Analysis and error evaluation for the clustered gradient

In order to analyze errors in clustered gradient the computation of reference gradient and the gradient obtained from *divide-and-conquer* procedure (clustered gradient) with different set of parameters are required. Then QM gradients computed for all clusters are analyzed with respect to the reference. Technically this is done by the *qr*.*granalyse* (“gradient analyse”) tool that automatically collects previously calculated and stored gradients and sorts them according to *fragment* and *buffer* size, and takes the gradient with the largest *buffer* (or if available from *super-sphere*) as reference unless manually specified otherwise. For each gradient an analysis described in the following is done and a PDB file is written for visualization. For each atom *i* in the model a measure of the atomic-gradient error (here referred to as the weighted difference gradient *δ*_*i*_) is computed and written into the B-factor field of the PDB file. That allows an easy color-shaded visualization of *δ*_*i*_ using standard biomolecular viewers.

In the error statistic we follow the idea of a regularized error [56], where we consider relative errors for large gradient differences and absolute errors for the small ones with the goal to bring both at a comparable level. As the regularisation we have chosen to use the median over all atomic gradients, instead of an arbitrarily pre-defined value as in [56], because the magnitude of the energy gradients is generally unknown. This choice was tested by detailed inspection of all gradients and their errors for the 3ftL case confirming that this metric follows what could be intuitively considered as small and large errors.

Firstly, we define the nuclear gradient matrix as

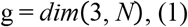

with *N* being the number of atoms. For each atom *i* the atomic difference gradient ΔG_*i*_ is expressed as the sum of the differences of the 3 cartesian component (c=x, y, z) with respect to the reference:

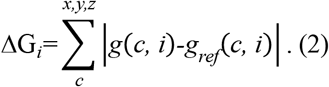

Finally, the weighted difference gradient *δ*_*i*_ for atom *i* is computed as

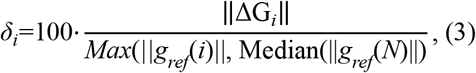

and the average over all *N* atoms is

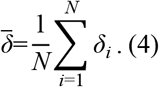

Where ||ΔG_*i*_|| is the norm of the difference gradient for the atom *i*, ||*g*_*ref*_(*i*)|| the norm of the reference gradient for the atom *i* and Median (||*g*_*ref*_(*N*)||) the median of all the reference gradient norms. Considering the reference gradient norm allows treating the magnitude of errors for small and large gradients on a more equal footing, while regularization with the Median gradient prevents too small gradients in the denominator from skewing the analysis. Moreover, for all cases where *δ*_*i*_ is larger than 100 (possible for example if the reference and tested gradients are of opposite sign) the final value is set to 100 to mimic the percentage and treat different cases on a comparable scale. Here we use a colour scheme with dark blue being 0 and 100 being red (in PyMol the command is “spectrum b, minimum=0, maximum=100”).

### 2.4 QM computations

The Q|R uses the ASE package [57] to interface with many modern QM computational software that can be used to calculate the gradients. In this work, we use TeraChem [42, 58, 59] and xtb [39] as QM calculators. TeraChem is a program employing Graphics Processing Units (GPUs) as a computing architecture for electronic structure calculations, which in turn allows efficient QM computations for thousands of atoms [58]. TeraChem calculations were performed with the Hartree-Fock (HF) method and the 6-31G basis set, with Grimme’s dispersion correction D3 [37] and the appropriate corrections for the basis set incompleteness via geometrical counterpoise model (gCP) [38]. The gCP and D3 corrections are implemented as part of *qr*.*refine* add-ons using standalone *gcp* [38] and *dftd3* [37] programs. Moreover, the environmental effects have been described employing the COSMO polarizable continuum solvent model [60]. The TeraChem computations have been performed on a Tesla K80 graphical card using 4 GPUs for the single QM run. The available computational resources consisted of total 48 cores, so that 12 single gradient computations can run at the same time.

The *xtb* computations were performed with the first-generation method named GFN1-xTB with bulk solvent effects treated using the internal generalized born (GB) solvation model. The GFN1-xTB calculations were executed on an Intel(R) Xeon(R) CPU E5-2690 v4 @ 2.10GHz CPU with a total of 28 cores or an Intel Core i9-10900X CPU with 3.7 GHz frequency and 20 cores.

## 3. Results

### 3.1 Sensitivity of the clustered gradient to the *super-sphere* size and QM methods

We start by comparing gradients computed for the whole model, without any clustering, but with increased surroundings, which is defined by the *super-sphere* R_ss_. We use gradients calculated with the largest *super-sphere* as the reference. The 3ftL peptide unit cell has only 209 atoms, but due to the close packing the number of atoms in the *super-sphere* increases rapidly with the Rss distance, at some point reaching as many atoms as in a relatively large model (such as 3q2c with 1611 atoms), as illustrated in Fig. 1a. The *super-sphere* obtained with the R_ss_ of 15 Å (7321 atoms for 3ftL and 8288 for 3q2c) is the largest for which the less expensive GFN1-xTB computations can be performed with our computer resources, which was considered as the reference. The trial gradients were computed with the R_ss_ sampled between 1 and 14 Å. Fig. 1b shows the average gradient error 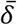. Starting from about 7 Å, the gradient errors become small and reach the plateau. Fig. 2 shows per-atom *δ*_*i*_ values for the 3ftL model calculated using four different Rss values. Clearly the choice of R_ss_ being 7 Å yields accurate gradients for all atoms (Fig. 2d). The R_ss_ of 7 Å is also largest for which more expensive HF-gCP-D3 computations can be performed. Fig. 1c shows that 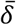 obtained at HF-gCP-D3 level are smaller than GFN1-xTB, albeit at a higher computational cost. Considering that the default radius of 10 Å would cause unnecessary computational effort due to significantly larger number of atoms (3220) the R_ss_ of 7 Å (1951 atoms) was used in subsequent analyses for 3ftL in conjunction with all QM methods. At variance, for 3q2c only GFN1-xTB method was considered so the default R_ss_ of 10 Å has been used as computationally feasible.

**Fig. 1.**
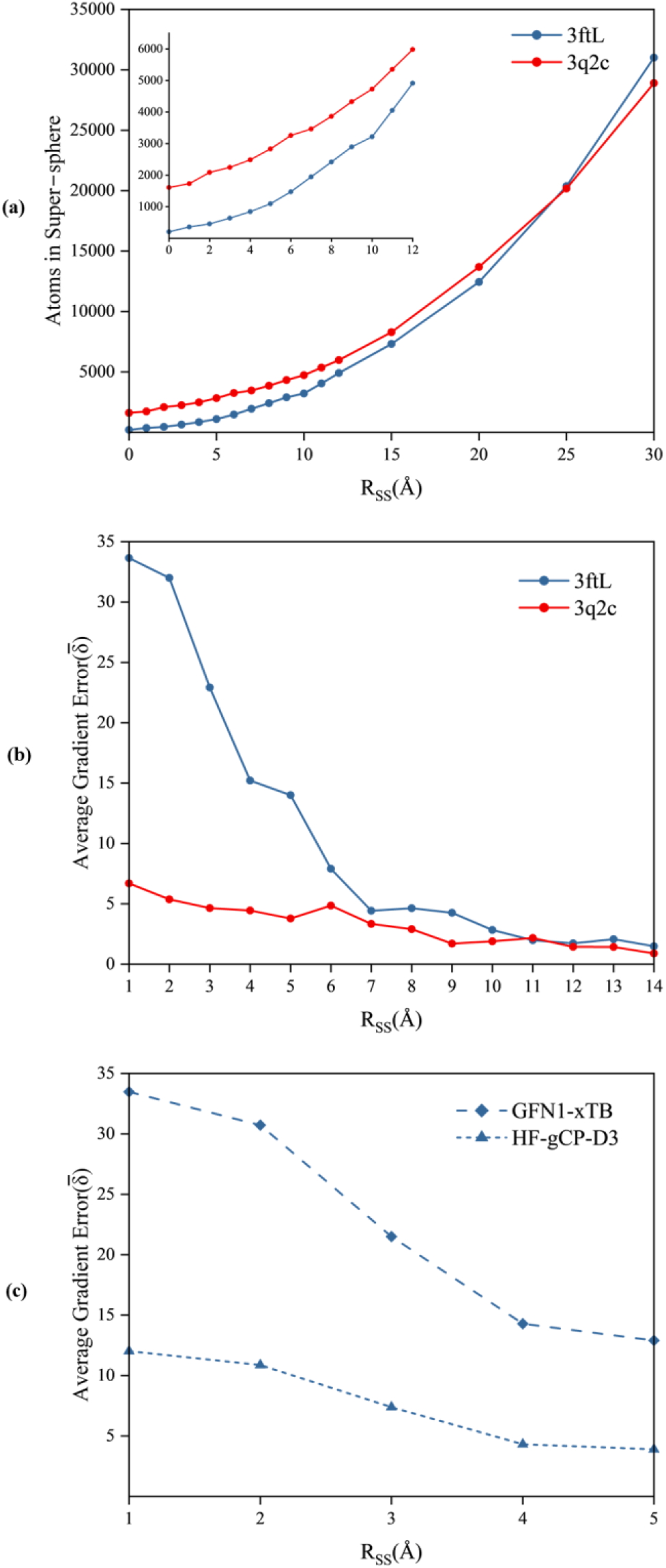
*Super-sphere* at different Rss (a) Number of atoms for 3ftL and 3q2c (the Rss range up to 12 Å is zoomed in the inset). Average atomic gradient error 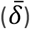 with respect to *super-sphere* Rss of (b) 15 Å for 3ftL and 3q2c computed at the GFN1-xTB level and (c) 7 Å for 3ftL computed at the HF-D3-gCP and GFN1-xTB levels.

**Fig. 2.**
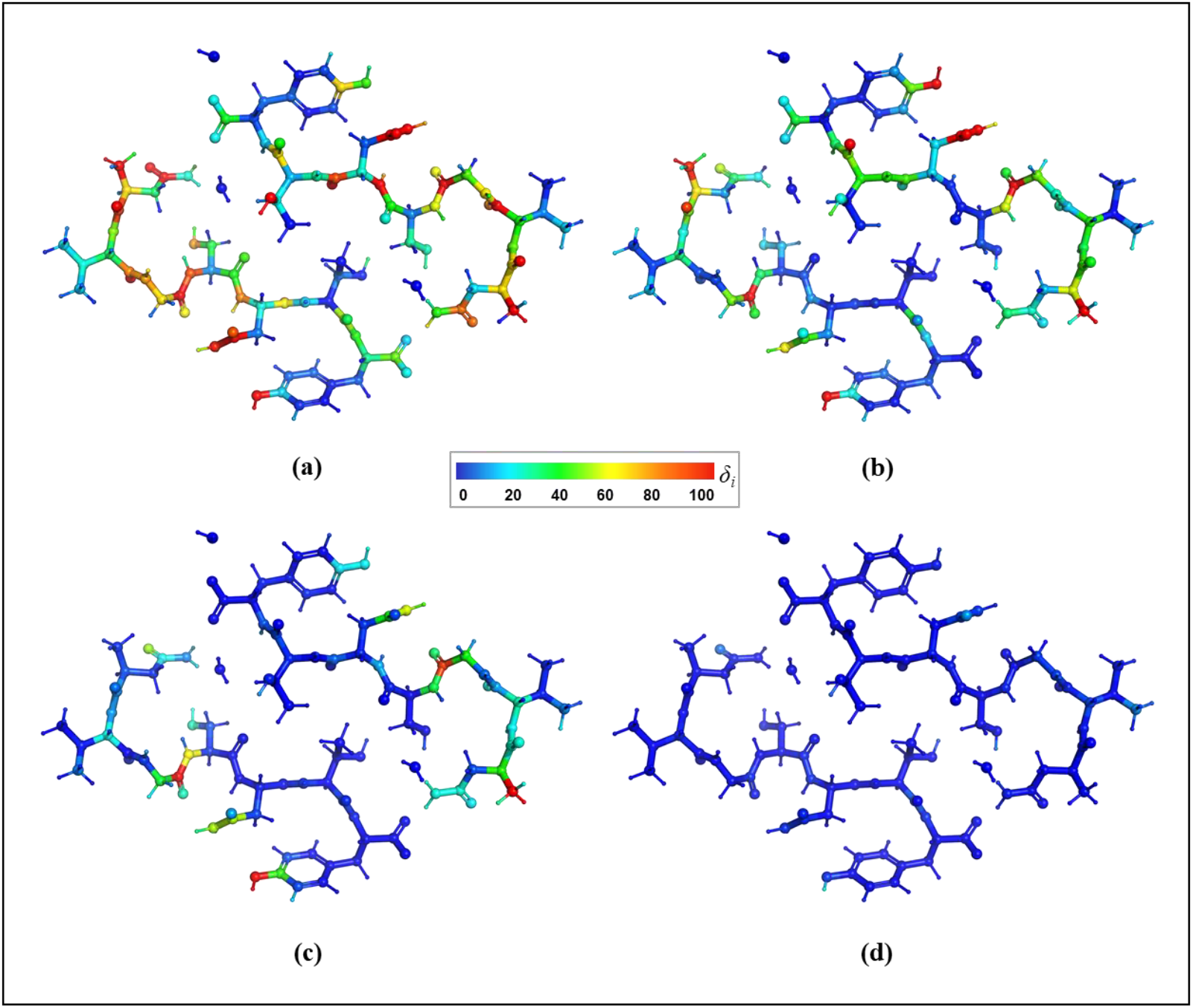
3ftL model colored by the atomic gradient errors for *super-sphere* with different R_ss_ (1 Å, 3 Å, 5 Å, 7 Å): (a, b, c, d) with respect to the reference with R_ss_=15 Å computed at the GFN1-xTB level.

### 3.2 Sensitivity of the clustered gradients to clustering

Here we analyze how the size and number of clusters affect the clustered gradient error. The 3ftL model can be partitioned into clusters using Q|R’s divide-and-conquer method in three different ways yielding 6, 3 and 2 clusters as shown in Fig. 3. For larger 3q2c protein there are more possibilities, so only some have been considered in this analysis. Fig. 4 shows average gradient error with respect to the clustering choice, in conjunction with smaller or larger *buffer* region. We observe that for given QM method and *buffering* the errors are very similar for all clustering choices.

**Fig. 3.**
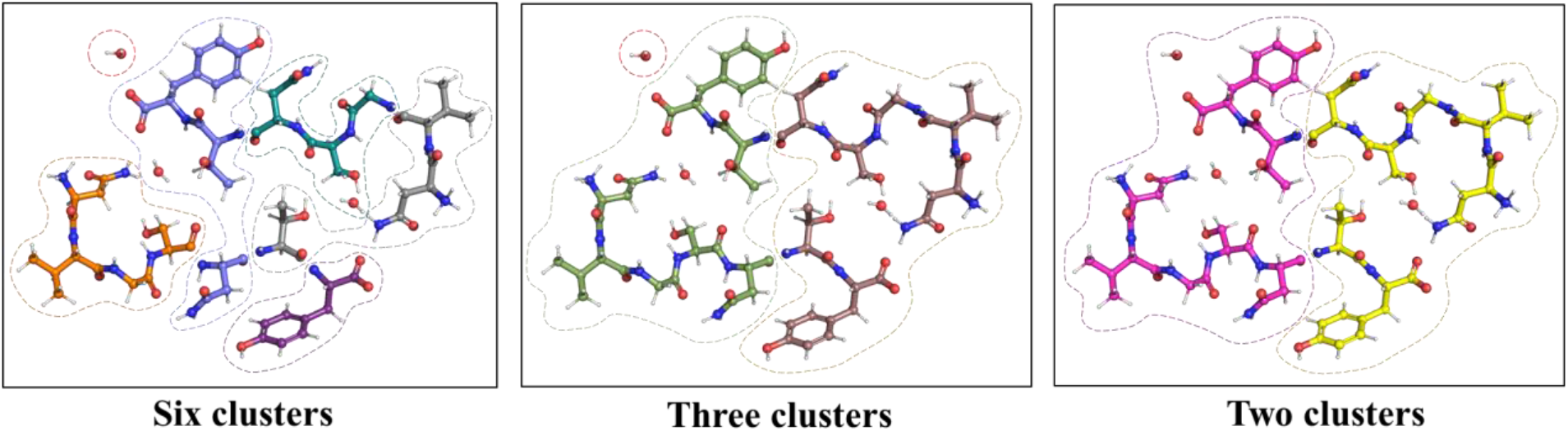
Three possible partitioning (clustering) of the 3ftL model

**Fig. 4.**
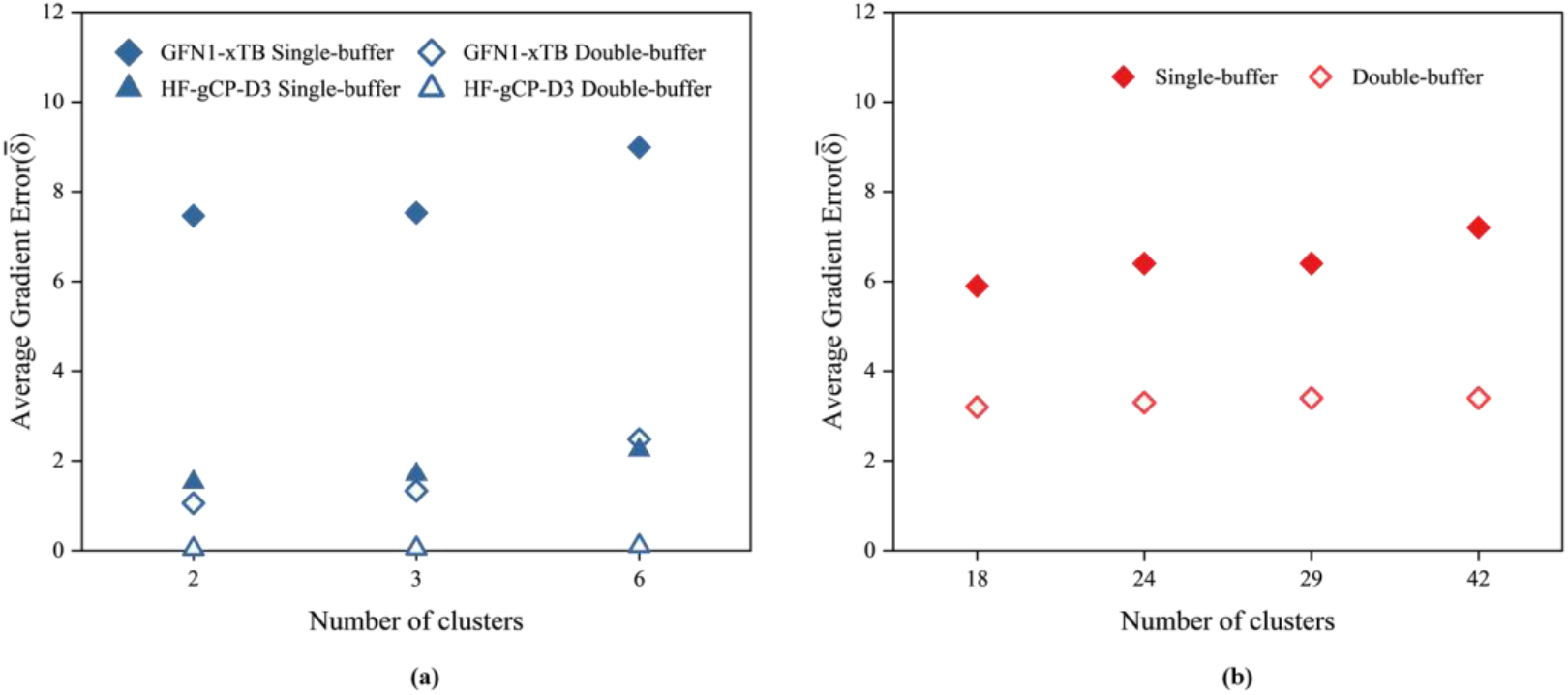
Average atomic gradient error for different clustering and *buffer*s (a) for 3ftL computed at the HF-gCP-D3 and GFN1-xTB levels. (b) for 3q2c computed at the GFN1-xTB level.

### 3.3 Sensitivity of the clustered gradients to *buffer* size

Here we study how the size of the *buffer* affects the accuracy of the clustered gradient. Clearly, the larger is the *buffer* the smaller are the errors in the clustered gradient (Fig. 4). For example, in case of 3ftL the largest errors resulting from using *double* versus *single-buffe*r are an order of magnitude smaller (1-2 versus 25). However, the larger the *buffe*r the longer it takes to calculate those gradients. Therefore, an optimal choice of the *buffer* is important. For that reason we analyse in more detail the atomic gradients obtained with different *buffer*s in order to rationalize situations leading to significant local errors. Highlighted cases include examples of local errors arising from cutting through covalent and non-covalent interactions within the model as well as due to expansion by the crystallographic symmetry.

#### 3.3.1 Covalent bonding within unit cell

The 3ftL peptide is composed of two symmetrical loops, each loop consisting of seven residues. In all clustering procedures, each of the A and B chains is cut at the N atom on Threonine (residue number 6), so the gradient error on these atoms has been analyzed in more detail.

When cutting the model into two clusters (Fig. 3), this N atom in chain B shows a negligible gradient error of 1.3 (Fig. 5a) but this error is 18.2 in the same atom in chain A (Fig. 5b). This can be explained by differences in the *buffer* region. In case of *single-buffer*, for the chain A only one residue is added after the cutting point (Fig. 5b), while for the chain B there are two residues past the cut (Fig. 5a). In the case of *double-buffer* both chains have at least three additional residues after the cutting point (Fig. 5c, d), which reduces the error from 18.2 and 1.3 (*single-buffer*) to 0.2 and 0.1.

**Fig. 5.**
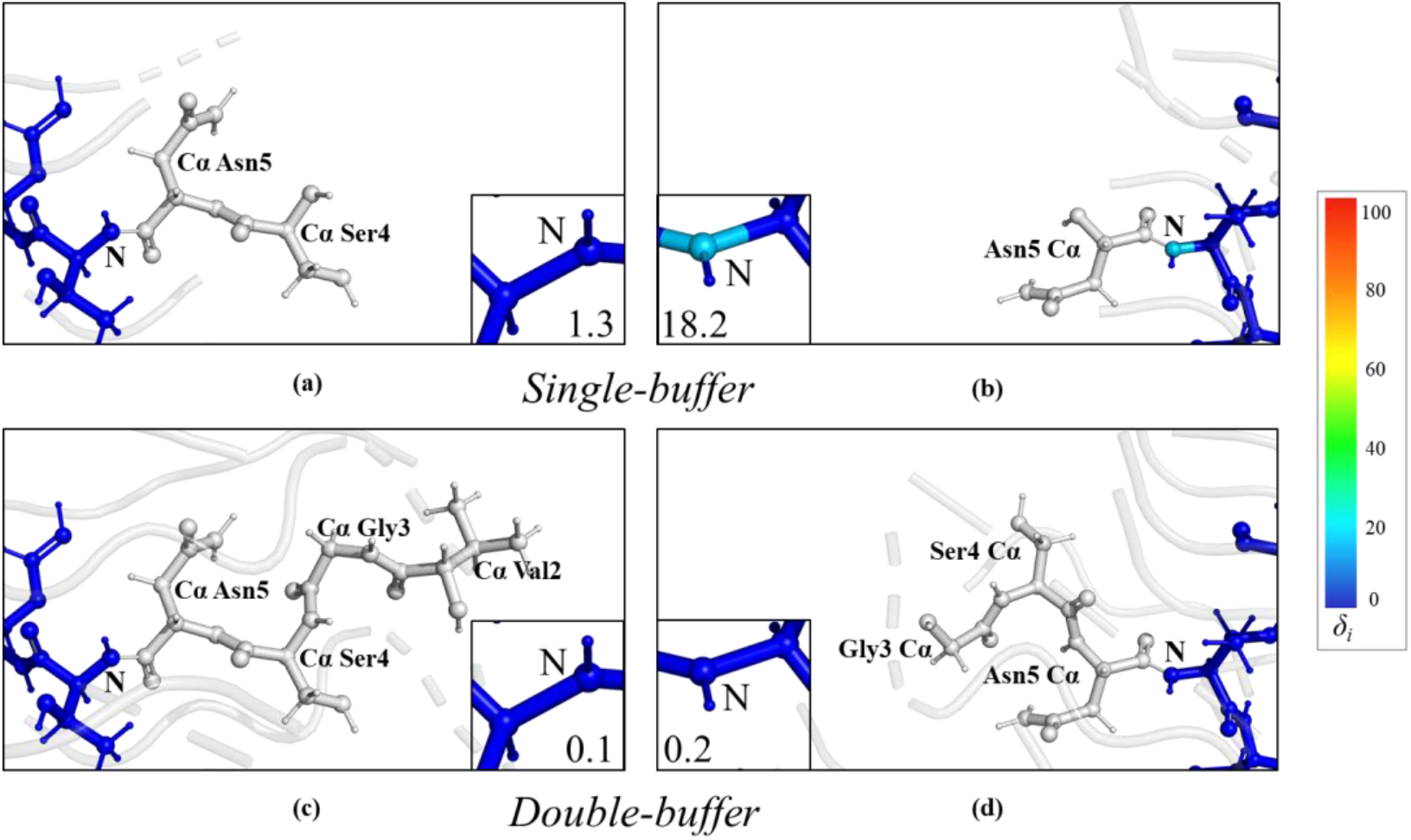
The 3ftL model divided into two clusters, showing fragments including *single-buffer* (a, b) and *double-buffer* (c, d). The *clusters* (colored by atomic *δ*_*i*_ values as computed at HF-gCP-D3 level) include two border-line nitrogen atoms. All atoms from *buffer* region are shown in grey, among them the residues directly behind the N atom in the chain are highlighted (grey ball and stick). The *δ*_*i*_ on both nitrogen atoms are reported in insets.

#### 3.3.2 Non-covalent interaction within unit cell

For the 3q2c the largest gradient errors appear on the oxygen and carbon atoms of the Valine residue 22 in chain A, with *δ*_*i*_ of 92.8 and 56.0, respectively (see Fig. 6a). This oxygen atom is involved into the hydrogen bond O···H-N with the Isoleucine 62 in chain A from the *buffer* region. In turn, this Isoleucine residue is truncated in the *single-buffer* and capped with a hydrogen atom. Using the *double-buffer* adds residues prior and past Isoleucine 62 (Fig. 6b), creating a six-residues chain and reducing the gradient error down to 0.8 and 3.3 on O and C, correspondingly.

**Fig. 6.**
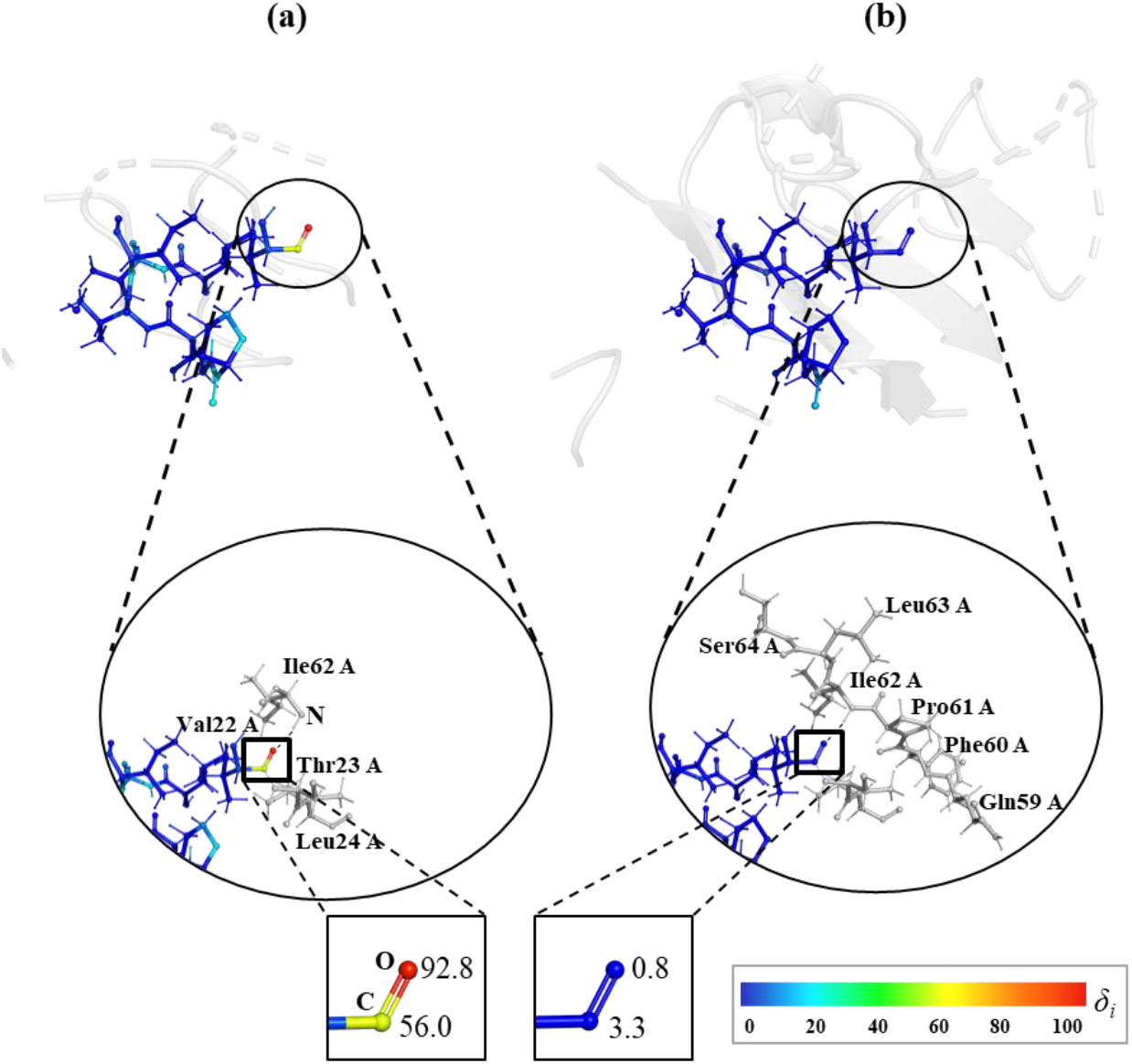
The 3q2c model, with non-covalent bonding inside unit cell, showing the *δ*_*i*_ on C and O atoms of residue Val 22 in chain A using (a) *single-* and (b) *double-buffer*.

#### 3.3.3 Interactions with symmetry copies

For the 3ftL model the largest errors in the clustered gradient were observed for H atoms involved in hydrogen bonding with symmetry copies. Namely, the hydrogen bonds D-H···A where either the hydrogen donor (D) or acceptor (A) belong to *buffer* region.

An example is the hydroxy groups in Tyrosine residue 7 in both chains that interact with symmetry copies (Fig. 7). Using the *single-buffer* results in errors around 11 and 23 on OH and HH in both chains, correspondingly, even though the directly interacting residues (Glycine 3 -- proton acceptor and Asparagine 5 -- proton donor) are present. Using the *double-buffer* includes more surroundings and that reduces these errors down to almost zero.

**Fig. 7.**
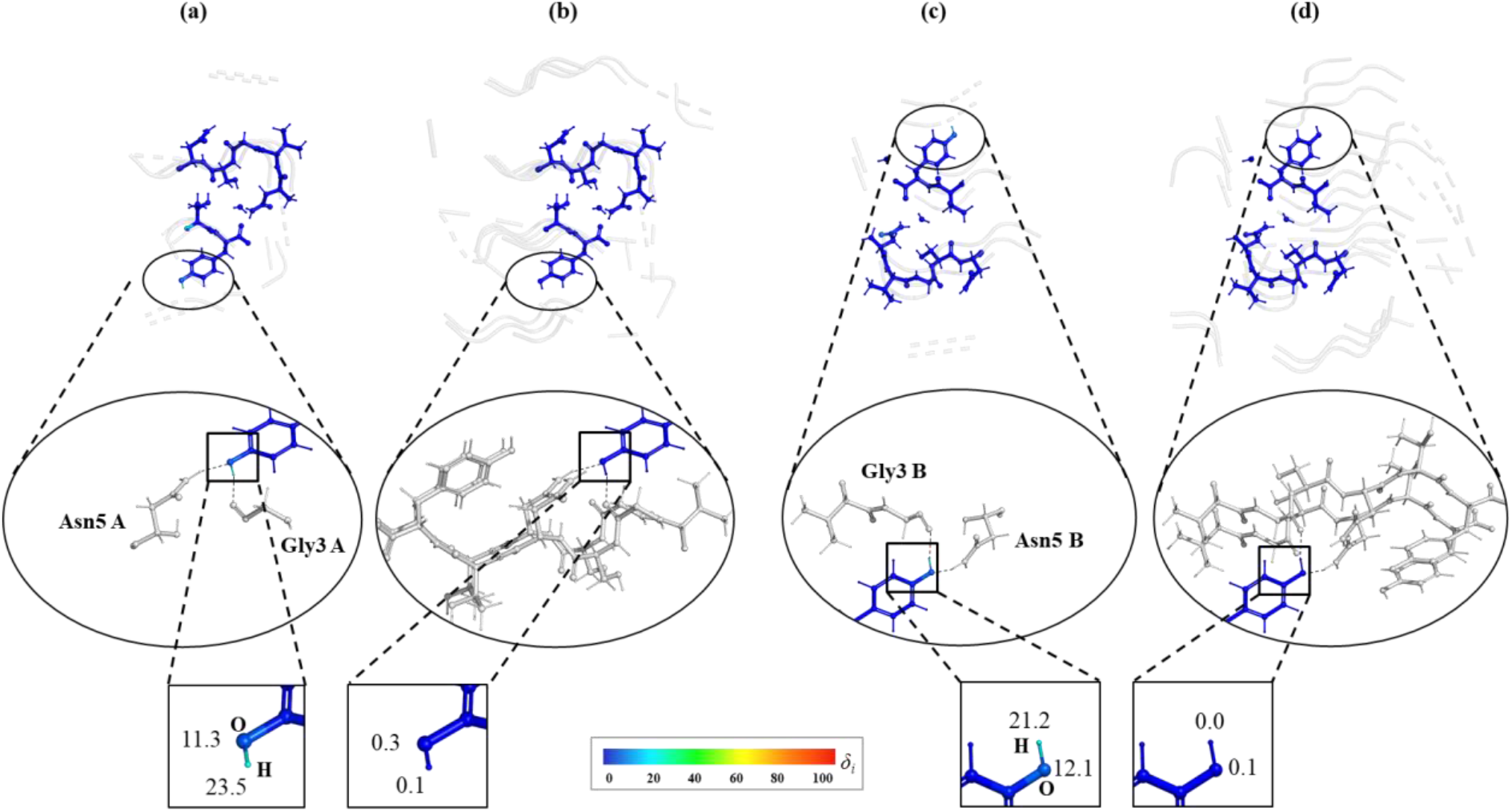
The interaction with symmetry copies for 3ftL model around H atom of TYR7. Different *buffer* environments for two clusters case (chain A a, b and chain B c, d): *single*-*buffer* (a, c) and *double-buffer* (b, d). All atoms in clusters are coloured by atomic *δ*_*i*_ values while the residues from *buffer* region are shown in grey.The enlarged circle in the middle highlights the hydrogen bonds formed with the atoms from the *buffer*. The boxes at the bottom zoom on O and H atoms from TYR7 and report their weighted difference gradient *δ*_*i*_ values.

**Fig. 8.**
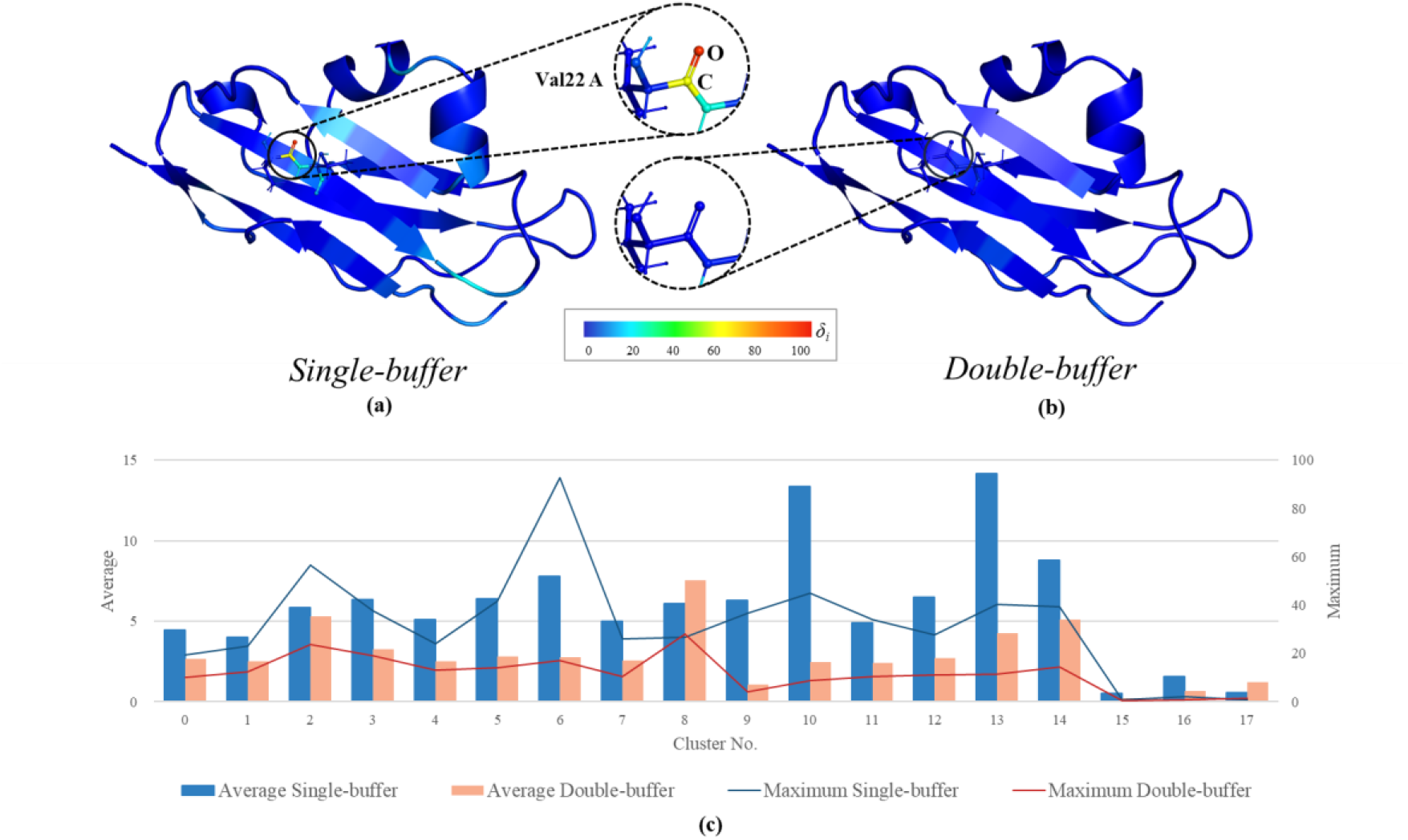
Atomic gradient errors for 3q2c protein: atomic *δ*_*i*_ in *single-* (a) and *double*-(b) *buffer* environments; average and maximum *δ*_*i*_ for each cluster (c).

### 3.4 Accuracy of clustered gradient for larger proteins

The gradient error analysis has been also performed for protein models with up to 7000 atoms (PDB codes: 3j63, 3a5x and chain C of 5fn5) that have shown significantly improved geometry metrics after quantum refinement in our recent study [51]. All gradients have been computed with the GFN1-xTB model and *single-buffer*, as employed in the Q|R #3 paper [51]. For the 3j63 and 5fn5 both *super-sphere* and *double-buffer* have been considered as reference, and only latter for the largest 3a5x. In all cases minimal gradient errors have been achieved with average and maximum *δ*_*i*_ below 1 and 10, respectively.

### 3.5 Timings

Among reliable *divide-and-conquer* schemes the choice of preferred one for computationally extensive refinement can be done based on the time required to compute single complete gradient. For that there are two limiting factors: (i) the size of the largest fragment 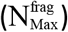 which defines the most time consuming QM computation; (ii) the total number of atoms in all fragments 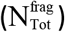 which defines the total CPU or GPU time needed to compute all gradients. We note that gradients for all fragments need to be computed in order to construct total clustered gradient. So, if all QM computations can run in parallel the size of largest fragment is the bottleneck factor, and division to larger number of smaller clusters is preferred. Second possibility is that all computations are run in serial. In that case division to smaller number of larger clusters, with smaller 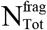 can be preferred.

However, in practice most common can be the third situation, that is number of clusters being larger than number of jobs which can be run in parallel. In this latter case, it might be cumbersome to decide *ad hoc* an optimal computational set-up, due to vast possibility of tunable parameters (clustering scheme as well as how many jobs (J) shall be run in parallel on how many cores (C) given the total number (M) of the latter (either CPU or GPU, M=J x C). These three situations are illustrated for 3ftL and 3q2c in Table 1 showing the timings required to compute a single gradient using HF-gCP-D3 (TeraChem) or GFN1-xTB (xtb).

**Table 1.** Number of atoms in *clusters* and *fragments* for 3ftL and 3q2c from different clustering choices along with the time needed to compute single complete gradient from TeraChem (HF-gCP-D3) or xtb (GFN1-xTB).

| Single-buffer |  |  |  |  |  | Double-buffer |  |  |  |
| --- | --- | --- | --- | --- | --- | --- | --- | --- | --- |
| Clust | $N_{\text{Max}}^{\text{clust}}$ | $N_{\text{Max}}^{\text{frag}}$ | $N_{\text{Tot}}^{\text{frag}}$ | $T^{\text{TC}}$ | $T^{\text{xtb}}$ | $N_{\text{Max}}^{\text{frag}}$ | $N_{\text{Tot}}^{\text{frag}}$ | $T^{\text{TC}}$ | $T^{\text{xtb}}$ |
| 3ftL |  |  |  |  |  |  |  |  |  |
| 6 | 50 | 563 | 2231 | 771 | 297 | 1560 | 6050 | 4975 | 1029 |
| 3 | 103 | 744 | 1564 | 1109 | 198 | 1630 | 3687 | 5167 | 943 |
| 2 | 106 | 787 | 1480 | 1161 | 204 | 1678 | 3203 | 5470 | 977 |
| 3q2c |  |  |  |  |  |  |  |  |  |
| 42 | 89 | 566 | 10405 |  | 740 | 1532 | 34092 |  | 5189 |
| 29 | 157 | 629 | 8209 |  | 575 | 1601 | 24933 |  | 4388 |
| 24 | 175 | 715 | 7345 |  | 518 | 1703 | 23024 |  | 4272 |
| 18 | 203 | 734 | 6501 |  | 453 | 1703 | 18821 |  | 4007 |

$N_{\text{Max}}^{\text{clust}}$ : Number of atoms in largest cluster.
$N_{\text{Max}}^{\text{frag}}$ : Number of atoms in largest fragment.
$N_{\text{Tot}}^{\text{frag}}$ : Total number of atoms in all fragments.
$T^{\text{TC}}$ : The computational time for gradient from TeraChem (TC) in seconds.
$T^{\text{xtb}}$ : The computational time for gradient from xtb in seconds.

For 3ftL the *cluster* size ranges from 3 (a single water molecule) to 106 atoms (one complete chain), with *fragment* sizes varying from 127 to 787 atoms for a *single-buffe*r and 532 to 1678 atoms for a *double-buffer*. This clearly indicates a notably higher computational cost for the latter. First situation is indicated by TeraChem computations, i.e. our GPU resources allowed computing gradients for all *fragment*s at once using HF-D3-gCP, so in practice the total timing was determined by 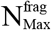. In the present case six *clusters* with the smallest possible *fragments* was the most effective scenario for both *single-* and *double-buffer*. Second situation is illustrated by GFN1-xTB computations (set to run sequentially) where two/three *clusters* is more favorable because the total number of atoms 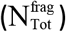 is reduced by about 30% compared to six *clusters*.

For 3q2c we consider only four possible clustering scenarios with the smallest number of clusters being 18. These computations were executed on the architecture with 20 cores, and we arbitrarily chosen to run 4 single gradient computations each one performed using 5 CPUs. This mimics the third situation: number of separate gradient calculations which run at a time is smaller than the total number of *clusters*. Therefore, computations are performed both in parallel and sequentially, which for this test-case leads to optimal situation with 18 clusters for both *single-* and *double-buffer*.

In more general terms the *gtest* procedure allows testing of realistic timings to find the most optimal for the specific model and computer resources. As during the course of a refinement many gradients will need to be calculated, the effort put into testing the timing of different *clustering* approaches will pay off in the long run.

### 3.5. Optimize time and gradient error

Increasing the *buffer* region clearly leads to improved atomic gradients. For 3q2c significantly lower *δ*_*i*_ are obtained for the *double*-*buffer*, with the all-atoms average and maximum of 3.2 and 28.1, respectively. For the *single-buffer* environment most of the atomic gradients also show small errors, but outliers with *δ*_*i*_ over 30 are present in half of the *clusters*, and over 50 in two *clusters*, leading to the *δ*_*i*_ averaged over all atoms equal to 5.9.

This shows that the extension of the *buffer* region for specific clusters represents a promising and cost-effective strategy to deal with well-localized outliers, avoiding an unnecessary *buffer* extension in well-behaved parts. Automatic gradient errors analysis allows implementing an adaptive “mixed” *buffer* strategy, into the *qr*.*refine*. The procedure starts from the computation of atomic *δ*_*i*_, which indicate if there are any clusters for which *single*-*buffer* environment is not sufficient. If that is the case, the *buffer* region is extended only for the specific *clusters*. This way the total number of atoms for which QM computations are required is reduced compared to the *double-buffering* applied to the whole model, saving computational resources. The number of atoms in all *fragments* obtained with different *buffering* strategies, relevant timings and maximum and average *δ*_*i*_ are reported for 3q2c in the Table 2. Considering a conservative maximum *δ*_*i*_ of 30 as the threshold, the total number of atoms in all *fragments* doubles with respect to *single-buffer*, but is still 30% smaller than for the full *double-buffer*. For the *δ*_*i*_ threshold of 50 there is an increase of atoms by 40% with respect to single, and decrease by 50% with respect to *double-buffer*, respectively. Corresponding computational time increases with respect to *single-buff*er by 5 to 2.5 times for Max>30 and Max>50 respectively, but is still about halved compared to *double-buffer*, without introducing outliers. As more experience is gained with other systems, the threshold can be fine-tuned in the future.

**Table 2.** The number of atoms in *fragments* from different *divide-and-conquer* schemes for 3q2c along with the time needed to compute single complete gradient from GFN1-xTB restraints.

| *Buffer* | Max number of atoms<br>in fragments | Total number of atoms<br>in all fragments | Time (s) | Average $\delta_i$ | Max $\delta_i$ |
| --- | --- | --- | --- | --- | --- |
| *Single* | 734 | 6501 | 453 | 5.9 | 92.8 |
| *Double* | 1703 | 18821 | 4007 | 3.2 | 28.1 |
| Mixed (Max>50) | 1703 | 9104 | 1112 | 5.5 | 45.1 |
| Mixed (Max>30) | 1703 | 13076 | 2393 | 4.1 | 27.8 |

Another possibility for reducing gradient errors is extending *buffer* for specific cluster by expanding *single-buffer* with the smallest possible number of residues. Fig. 9 shows *δ*_*i*_ on atoms involved in hydrogen bonds with buffer region, where 3q2c Val22 acts as a proton donor (N-H···O) or acceptor (C=O···H-N). The latter correspond to the largest gradient errors for *single-buffer* as discussed in section 3.2.2. By adding into buffer just one residue (Pro61) large errors on oxygen and carbon atoms are reduced from *δ*_*i*_ of 92.8 and 56.0, respectively to about 25 on both atoms (Fig. 9a). However, at the same time errors on nitrogen and hydrogen from adjacent hydrogen bond increase with respect to *single-buffer* case, from 8.5 and 12.2 to 29.7 and 61.7, respectively. So, another unbalanced environment is created. The latter problem is resolved by adding one residue after Ile62 (Fig. 9b), which reduces errors below 20 on all N, H, C and O of Val22. Overall, a balanced description requires symmetric buffering – one (Fig. 9b), or two (Fig. 9d) residues added on both sides of Ile62, while asymmetric situations (Fig. 9a, c) show larger errors. This analysis indicates that definition of reduced, yet smaller buffer region might be a non-trivial task.

**Fig. 9.**
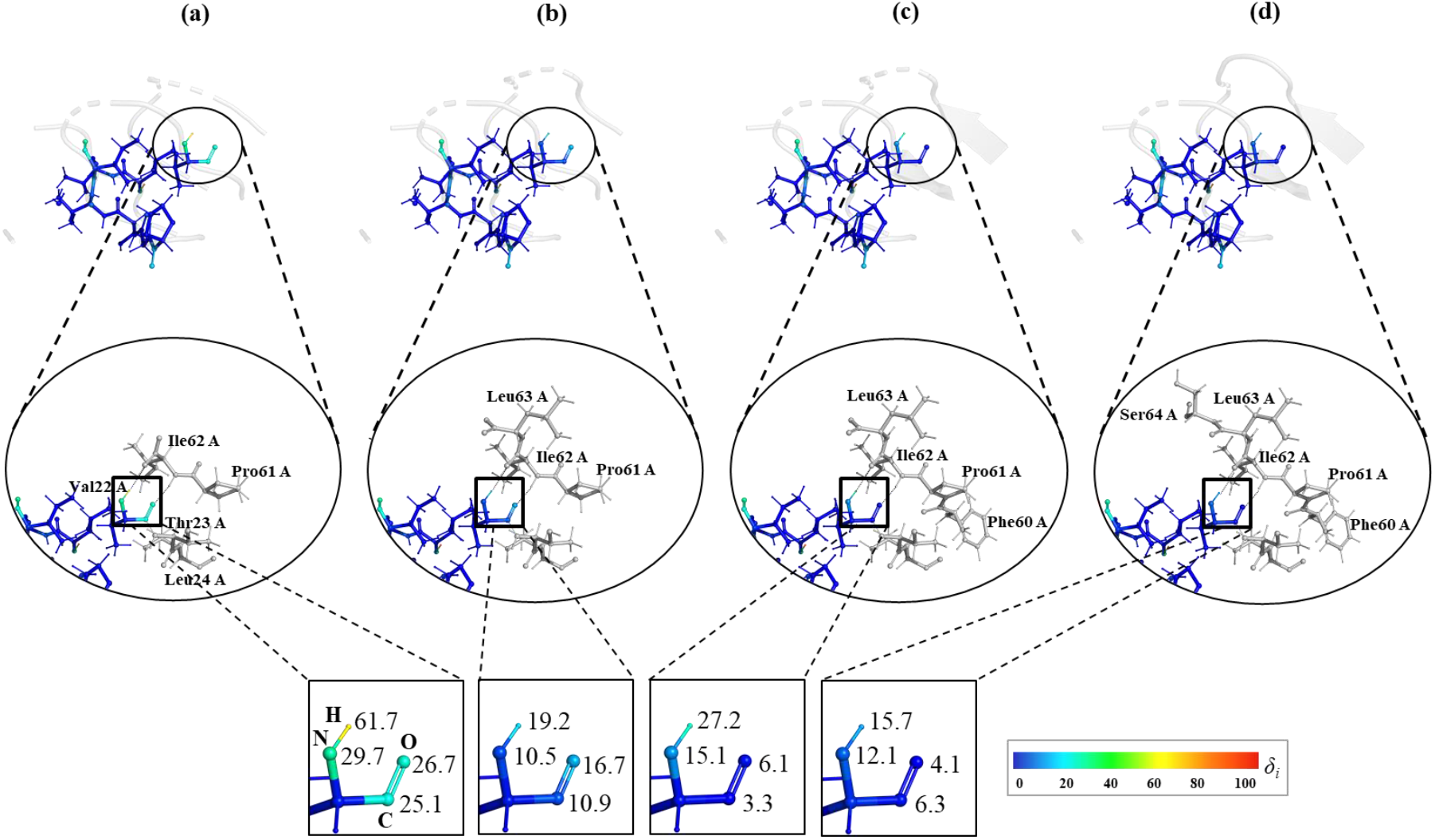
The 3q2c model, *δ*_*i*_ on C, O, N and H atoms of residue Val 22 in chain A for *buffer* cases between *single-* and *double-buffer* (reported in Fig. 6a and 6b, respectively). Possibilities consider adding one-by-one residues to the amino acid backbone in the fragment.

## Conclusions

Herein we analyze the robustness of the QM nuclear gradient from the *divide-and-conquer* procedure employed during quantum refinement in the Q|R project. Moreover, we present new parts of the Q|R code called *gtest* and *granalyze* that help to automate assessment of the gradient quality.

The procedure was showcased for the 3ftL peptide and 3q2c protein where errors in energy gradients were analysed with respect to *super-sphere* size, *clusters* of different sizes and increasing *buffer* regions, by comparing to a reference gradient obtained from a *super-sphere* calculation with large R_ss_. We introduced an error metric that allows to pin-point errors in the gradient to specific atoms or groups that then can be checked for missing interaction with, e.g. symmetry-copies or near-by residues, due to an insufficient *buffer* region. To simplify the analysis for the user the tool writes PDBs files with color coding to allow an intuitive visual inspection.

Different QM methods show a similar sensitivity to the clustering parameters, with overall gradient errors for using HF-gCP-D3/6-31G, being smaller than for the semi-empirical quantum methods GFN1-xTB.

Moreover, we showed that by applying the *double-buffer* procedure essentially error-free gradients can be obtained for the case when the *super-sphere* calculation is computationally prohibitive.

The presented *granalyze* procedure is an effective tool by which diverse *divide-and-conquer* schemes, varying by number of residues in the *cluster* or the applied *buffer* region can be easily tested, the most computationally efficient setup chosen and their errors analyzed. This allows also implementing new strategies for the efficient choice of adaptive *buffer* region during quantum refinement, avoiding unnecessary environment expansion for the parts of molecular model with already error-free gradient. Finally, both automatic partitioning scheme and gradient analysis tools from Q|R can be employed to define d*ivide-and-conquer* strategies for other QM computations of bio-macromolecules, beyond the quantum refinement.

All the tools developed within the Q|R project are available at https://github.com/qrefine/qrefine.

Gradient error analysis results and related data are available at: https://github.com/qrefine/QR4-Gtest.

## Acknowledgments

MB and YW acknowledge the financial support from the National Natural Science Foundation of China (Grant No. 31870738). MB also acknowledges support of the COST Action CA21101 “COSY”.

## Author Contributions Statement

MB, HK and PVA wrote the main manuscript text. HK developed tools describe in the manuscript. YW performed all computations, analysis and prepared figures. PVA, NWM, MPW and HK developed the overall code. All authors reviewed the manuscript.

## References

1. Shi Y (2014) A glimpse of structural biology through X-ray crystallography. Cell 159:995–1014. https://doi.org/10.1016/j.cell.2014.10.051

2. Branden CI, Tooze J (2012) Introduction to protein structure, Garland Science, New York. https://doi.org/10.1201/9781136969898

3. Borbulevych OY, Plumley JA, Martin RI et al (2014) Accurate macromolecular crystallographic refinement: incorporation of the linear scaling, semiempirical quantum-mechanics program DivCon into the PHENIX refinement package. Acta Crystallogr Sect D Biol Crystallogr 70:1233–1247. https://doi.org/10.1107/S13990047140022604.

4. Senthil R, Sakthivel M, Usha S (2021) Structure-based drug design of peroxisome proliferator-activated receptor gamma inhibitors: ferulic acid and derivatives. J Biomol Struct Dyn 39:1295–1311. https://doi.org/10.1080/07391102.2020.1740790

5. Kordbacheh S, Kasko AM (2021) Peptide and protein engineering by modification of backbone and sidechain functional groups. Polym Int 70:889–896. https://doi.org/10.1002/pi.6208

6. Urzhumtsev AG, Lunin VY (2019) Introduction to crystallographic refinement of macromolecular atomic models. Crystallogr Rev 25:164–262. https://doi.org/10.1080/0889311X.2019.1631817

7. Waser J (1963) Least-squares refinement with subsidiary conditions. Acta Cryst 16:1091–1094. https://doi.org/10.1107/S0365110X63002929

8. Engh R, Huber R (2001) International Tables for Crystallography, Vol. F, edited by MG Rossmann & E. Arnold, Dordrecht: Kluwer Academic Publishers 382–392

9. Groom CR, Bruno IJ, Lightfoot MP et al (2016) The Cambridge structural database. Acta Crystallogr B Struct Sci Cryst Eng Mater 72:171–179. https://doi.org/10.1107/S2052520616003954

10. Taylor R, Wood PA (2019) A million crystal structures: The whole is greater than the sum of its parts. Chem Rev 119:9427–9477. https://doi.org/10.1021/acs.chemrev.9b00155

11. Vaitkus A, Merkys A, Gražulis S (2021) Validation of the crystallography open database using the crystallographic information framework. J Appl Crystallogr 54:661–672. https://doi.org/10.1107/S1600576720016532

12. Berkholz DS, Shapovalov MV, Dunbrack Jr RL et al (2009) Conformation dependence of backbone geometry in proteins. Structure 17:1316–1325. https://doi.org/10.1016/j.str.2009.08.012

13. Moriarty NW, Tronrud DE, Adams PD et al (2014) Conformation-dependent backbone geometry restraints set a new standard for protein crystallographic refinement. FEBS J 281:4061–4071. https://doi.org/10.1111/febs.12860

14. Moriarty NW, Tronrud DE, Adams PD et al (2016) A new default restraint library for the protein backbone in Phenix: a conformation-dependent geometry goes mainstream. Acta Crystallogr D Struct Biol 72:176–179. https://doi.org/10.1107/S2059798315022408

15. Liebeschuetz J, Hennemann J, Olsson T et al (2012) The good, the bad and the twisted: a survey of ligand geometry in protein crystal structures. J Comput Aided Mol Des 26:169–183. https://doi.org/10.1007/s10822-011-9538-6

16. Janowski PA, Moriarty NW, Kelley BP et al (2016) Improved ligand geometries in crystallographic refinement using AFITT in PHENIX. Acta Crystallogr D Struct Biol 72:1062–1072. https://doi.org/10.1107/S2059798316012225

17. Peach ML, Cachau RE, Nicklaus MC (2017) Conformational energy range of ligands in protein crystal structures: the difficult quest for accurate understanding. J Mol Recognit 30:e2618. https://doi.org/10.1002/jmr.2618

18. Liebeschuetz JW (2021) The Good, the Bad, and the Twisted Revisited: An Analysis of Ligand Geometry in Highly Resolved Protein–Ligand X-ray Structures. J Med Chem 64:7533–7543. https://doi.org/10.1021/acs.jmedchem.1c00228

19. Brereton AE, Karplus PA (2015) Native proteins trap high-energy transit conformations. Sci Adv 1:e1501188. https://doi.org/10.1126/sciadv.1501188

20. Jiang Z, Biczysko M, Moriarty NW (2018) Accurate geometries for “Mountain pass” regions of the Ramachandran plot using quantum chemical calculations. Proteins 86:273–278. https://doi.org/10.1002/prot.25451

21. Moriarty NW, Liebschner D, Tronrud DE et al (2020) Arginine off-kilter: guanidinium is not as planar as restraints denote. Acta Crystallogr D Struct Biol 76:1159–1166. https://doi.org/10.1107/S2059798320013534

22. Qi HW, Kulik HJ (2019) Evaluating unexpectedly short non-covalent distances in x-ray crystal structures of proteins with electronic structure analysis. J Chem Inf Model 59:2199–2211. https://doi.org/10.1021/acs.jcim.9b00144

23. Moriarty NW, Janowski PA, Swails JM et al (2020) Improved chemistry restraints for crystallographic refinement by integrating the Amber force field into Phenix. Acta Crystallogr D Struct Biol 76:51–62. https://doi.org/10.1107/S2059798319015134

24. Borbulevych O, Martin RI, Westerhoff LM (2018) High-throughput quantum-mechanics/molecular-mechanics (ONIOM) macromolecular crystallographic refinement with PHENIX/DivCon: the impact of mixed Hamiltonian methods on ligand and protein structure. Acta Crystallogr D Struct Biol 74:1063–1077. https://doi.org/10.1107/S2059798318012913

25. Genoni A, Bučinský L, Claiser N et al (2018) Quantum crystallography: Current developments and future perspectives. Chem Eur J 24:10881–10905. https://doi.org/10.1002/chem.201705952

26. Caldararu O, Manzoni F, Oksanen E et al (2019) Refinement of protein structures using a combination of quantum-mechanical calculations with neutron and X-ray crystallographic data. Acta Crystallogr D Struct Biol 75:368–380. https://doi.org/10.1107/S205979831900175X

27. Yan Z, Li X, Chung LW (2021) Multiscale Quantum Refinement Approaches for Metalloproteins. J Chem Theory Comput 17:3783–3796. https://doi.org/10.1021/acs.jctc.1c00148

28. Bergmann J, Oksanen E, Ryde U (2022) Combining crystallography with quantum mechanics. Curr Opin Struct Biol 72:18–26. https://doi.org/10.1016/j.sbi.2021.07.002

29. Merz Jr KM (2014) Using quantum mechanical approaches to study biological systems. Acc Chem Res 47:2804–2811. https://doi.org/10.1021/ar5001023

30. Zheng M, Reimers JR, Waller MP et al (2017) Q| R: quantum-based refinement. Acta Crystallogr D Struct Biol 73:45–52. https://doi.org/10.1107/S2059798316019847

31. Liebschner D, Afonine PV, Baker ML et al (2019) Macromolecular structure determination using X-rays, neutrons and electrons: recent developments in Phenix. Acta Crystallogr D Struct Biol 75:861–877. https://doi.org/10.1107/S2059798319011471

32. Vagin AA, Murshudov GN (2004) IUCr Comput. Comm. Newsl. 4:59–72.

33. Vagin AA, Steiner RA, Lebedev AA et al (2004) REFMAC5 dictionary: Organization of prior chemical knowledge and guidelines for its use. Acta Crystallogr Sect D Biol Crystallogr 60:2184–2195. https://doi.org/10.1107/S0907444904023510

34. Senn HM, Thiel W (2009) QM/MM methods for biomolecular systems. Angew Chem Int Ed 48:1198–1229. https://doi.org/10.1002/anie.200802019

35. Ryde U (2016) QM/MM calculations on proteins. Meth Enzymol 577:119–158. https://doi.org/10.1016/bs.mie.2016.05.014

36. Canfield P, Dahlbom MG, Hush NS et al (2006) Density-functional geometry optimization of the 150 000-atom photosystem-I trimer. J Chem Phys 124:024301. https://doi.org/10.1063/1.2148956

37. Grimme S, Antony J, Ehrlich S et al (2010) A consistent and accurate ab initio parametrization of density functional dispersion correction (DFT-D) for the 94 elements H-Pu. J Chem Phys 132:154104. https://doi.org/10.1063/1.3382344

38. Kruse H, Grimme S (2012) A geometrical correction for the inter-and intra-molecular basis set superposition error in Hartree-Fock and density functional theory calculations for large systems. J Chem Phys 136:04B613. https://doi.org/10.1063/1.3700154

39. Grimme S, Bannwarth C, Shushkov P (2017) A robust and accurate tight-binding quantum chemical method for structures, vibrational frequencies, and noncovalent interactions of large molecular systems parametrized for all spd-block elements (Z= 1–86). J Chem Theory Comput 13:1989–2009. https://doi.org/10.1021/acs.jctc.7b00118

40. Klamt A, Schüürmann G (1993) COSMO: a new approach to dielectric screening in solvents with explicit expressions for the screening energy and its gradient. J Chem Soc, Perkin Trans 2 799–805. https://doi.org/10.1039/P29930000799

41. Carlsen M, Røgen P (2015) Protein structure refinement by optimization. Proteins 83:1616–1624. https://doi.org/10.1002/prot.24846

42. Titov AV, Ufimtsev IS, Luehr N et al (2013) Generating efficient quantum chemistry codes for novel architectures. J Chem Theory Comput 9:213–221. https://doi.org/10.1021/ct300321a

43. Herbert JM (2019) Fantasy versus reality in fragment-based quantum chemistry. J Chem Phys 151:170901. https://doi.org/10.1063/1.5126216

44. Gordon MS, Fedorov DG, Pruitt SR et al (2012) Fragmentation methods: A route to accurate calculations on large systems. Chem Rev 112:632–672. https://doi.org/10.1021/cr200093j

45. Collins MA, Bettens RP (2015) Energy-based molecular fragmentation methods. Chem Rev 115:5607–5642. https://doi.org/10.1021/cr500455b

46. Raghavachari K, Saha A (2015) Accurate composite and fragment-based quantum chemical models for large molecules. Chem Rev 115:5643–5677. https://doi.org/10.1021/cr500606e

47. Liu J, He X (2020) Fragment-based quantum mechanical approach to biomolecules, molecular clusters, molecular crystals and liquids. Phys Chem Chem Phys 22:12341–12367. https://doi.org/10.1039/D0CP01095B

48. Kitaura K, Ikeo E, Asada T et al (1999) Fragment molecular orbital method: an approximate computational method for large molecules. Chem Phys Lett 313:701–706. https://doi.org/10.1016/S0009-2614(99)00874-X

49. Zheng M, Moriarty NW, Xu Y et al (2017) Solving the scalability issue in quantum-based refinement: Q| R# 1. Acta Crystallogr D Struct Biol 73:1020–1028. https://doi.org/10.1107/S2059798317016746

50. Zheng M, Biczysko M, Xu Y et al (2020) Including crystallographic symmetry in quantum-based refinement: Q| R# 2. Acta Crystallogr D Struct Biol 76:41–50. https://doi.org/10.1107/S2059798319015122

51. Wang L, Kruse H, Sobolev OV et al (2020) Real-space quantum-based refinement for cryo-EM: Q| R# 3. Acta Crystallogr D Struct Biol 76:1184–1191. https://doi.org/10.1107/S2059798320013194

52. Schmitz S, Seibert J, Ostermeir K et al (2020) Quantum chemical calculation of molecular and periodic peptide and protein structures. J Phys Chem B 124:3636–3646. https://doi.org/10.1021/acs.jpcb.0c00549

53. Riek R (2017) The three-dimensional structures of amyloids. Cold Spring Harb Perspect Biol 9:a023572. https://doi.org/10.1101/cshperspect.a023572

54. Burley SK, Berman HM, Bhikadiya C et al (2019) Protein Data Bank: the single global archive for 3D macromolecular structure data. Nucleic Acids Res 47:D520–D528. https://doi.org/10.1093/nar/gky949

55. Afonine PV, Grosse-Kunstleve RW, Echols N et al (2012) Towards automated crystallographic structure refinement with phenix. refine. Acta Crystallogr Sect D Biol Crystallogr 68:352–367. https://doi.org/10.1107/S0907444912001308

56. Hait D, Head-Gordon M (2018) How accurate is density functional theory at predicting dipole moments? An assessment using a new database of 200 benchmark values. J Chem Theory Comput 14:1969–1981. https://doi.org/10.1021/acs.jctc.7b01252

57. Larsen AH, Mortensen JJ, Blomqvist J et al (2017) The atomic simulation environment—a Python library for working with atoms. J Phys: Condens Matter 29:273002. https://doi.org/10.1088/1361-648X/aa680e

58. Seritan S, Bannwarth C, Fales BS et al (2021) TeraChem: A graphical processing unit-accelerated electronic structure package for large-scale ab initio molecular dynamics. Wiley Interdiscip Rev Comput Mol Sci 11:e1494. https://doi.org/10.1002/wcms.1494

59. Ufimtsev IS, Martinez TJ (2009) Quantum chemistry on graphical processing units. 3. Analytical energy gradients, geometry optimization, and first principles molecular dynamics. J Chem Theory Comput 5:2619–2628. https://doi.org/10.1021/ct9003004

60. Liu F, Luehr N, Kulik HJ et al (2015) Quantum chemistry for solvated molecules on graphical processing units using polarizable continuum models. J Chem Theory Comput 11:3131–3144. https://doi.org/10.1021/acs.jctc.5b00370

